# Temporal dynamics of SARS-CoV-2 mutation accumulation within and across infected hosts

**DOI:** 10.1101/2021.01.19.427330

**Authors:** Andrew L. Valesano, Kalee E. Rumfelt, Derek E. Dimcheff, Christopher N. Blair, William J. Fitzsimmons, Joshua G. Petrie, Emily T. Martin, Adam S. Lauring

**Affiliations:** Division of Infectious Diseases, Department of Internal Medicine, University of Michigan, Ann Arbor, MI, USA; Department of Microbiology and Immunology, University of Michigan, Ann Arbor, MI, USA; Division of Hospital Medicine, Department of Internal Medicine, University of Michigan, Ann Arbor, MI, USA; Department of Epidemiology, University of Michigan, Ann Arbor, MI, USA

**Author notes:** Corresponding author Adam S. Lauring, 1150 W. Medical Center Dr., MSRB1 Room 5510B, Ann Arbor, MI 48109-5680.

**Keywords:** SARS-CoV-2, intrahost diversity, sequencing, transmission, evolution

## Abstract

Analysis of SARS-CoV-2 genetic diversity within infected hosts can provide insight into the generation and spread of new viral variants and may enable high resolution inference of transmission chains. However, little is known about temporal aspects of SARS-CoV-2 intrahost diversity and the extent to which shared diversity reflects convergent evolution as opposed to transmission linkage. Here we use high depth of coverage sequencing to identify within-host genetic variants in 325 specimens from hospitalized COVID-19 patients and infected employees at a single medical center. We validated our variant calling by sequencing defined RNA mixtures and identified a viral load threshold that minimizes false positives. By leveraging clinical metadata, we found that intrahost diversity is low and does not vary by time from symptom onset. This suggests that variants will only rarely rise to appreciable frequency prior to transmission. Although there was generally little shared variation across the sequenced cohort, we identified intrahost variants shared across individuals who were unlikely to be related by transmission. These variants did not precede a rise in frequency in global consensus genomes, suggesting that intrahost variants may have limited utility for predicting future lineages. These results provide important context for sequence-based inference in SARS-CoV-2 evolution and epidemiology.

## Introduction

Over the course of the SARS-CoV-2 pandemic, whole genome sequencing has been widely used to characterize patterns of broad geographic spread, transmission in local clusters, and the spread of specific viral variants^1–6^. Early reports demonstrated that SARS-CoV-2 exhibits genetic diversity within infected hosts, but this has been less studied than consensus-level genomic diversity^7^. Intrahost diversity is an important complement to consensus sequencing. Patterns of viral intrahost diversity throughout individual infections can suggest the relative importance of natural selection and stochastic genetic drift^8^. Shared intrahost variants between individuals can reveal loci under convergent evolution and enable measurement of the transmission bottleneck, a critical determining factor in the spread of new genetic variants^9,10^. Studies of SARS-CoV-2 intrahost diversity may shed light on selective pressures applied at the individual level, such as antivirals and antibody-based therapeutics. While a clear understanding of within-host evolution can inform how SARS-CoV-2 spreads on broader scales, there have been relatively few comprehensive studies of intrahost dynamics^9,11,12^.

Sequencing of intrahost populations can also potentially be applied to genomic epidemiology^13^. A common goal in sequencing specimens from case clusters is to infer transmission linkage, which can guide future public health and infection control interventions. However, the relatively low substitution rate and genetic diversity of SARS-CoV-2 present challenges to inference of individual transmission pairs^13,14^. In the pandemic setting, there is a non-negligible chance that two individuals who are epidemiologically unrelated could be infected with nearly identical viral genomes. Viruses from a single local outbreak may have few differentiating substitutions, limiting the ability of sequencing to resolve exact transmission chains. Identification of shared intrahost variants between individuals has been explored in other pathogens to overcome this obstacle^15–19^. However, use of this approach for SARS-CoV-2 will depend on a solid understanding of the forces that shape the generation and spread of genetic variants. There are several unresolved questions that will dictate the utility of intrahost diversity for genomic epidemiology. First, there must be sufficient intrahost diversity generated during acute infection prior to a transmission event. How much intrahost diversity is accumulated over time from infection onset is currently unknown. Second, the population bottleneck during transmission must be sufficiently wide to allow minor variants to be transmitted to recipient hosts^20,21^. Third, *de novo* generation of the same minor variants across multiple infections must be sufficiently rare. Independent generation of shared minor variants by positive selection or genetic drift in unrelated hosts could confound transmission inference^15^. Finally, measurements of intrahost diversity must be accurate and account for several potential sources of error^22,23^. Although previous studies have described within-host variation of SARS-CoV-2^7,9,11,12,24–26^, few have addressed the sources of systematic errors and batch effects in variant identification. To assess the utility of SARS-CoV-2 intrahost diversity for transmission inference, we need a clearer understanding of its temporal variation throughout infection and the extent of convergent evolution across individuals. Addressing these questions will also be valuable for understanding SARS-CoV-2 evolution.

Here, we sequenced SARS-CoV-2 genomes from 325 residual upper respiratory samples from hospitalized patients and employees at the University of Michigan. To validate our sequencing approach, we sequenced defined mixtures of two synthetic RNA controls and found that low input viral load decreases the specificity of variant calling. We find that observed intrahost diversity does not vary significantly by day since symptom onset. Intrahost variants can be shared between individuals that are unlikely to be related by transmission, suggesting that variants can arise by parallel evolution. These results inform our understanding of SARS-CoV-2 diversification in human hosts and highlight important considerations for sequence-based inference in the virus’s genomic epidemiology.

## Results

We retrieved respiratory specimens collected through diagnostic testing from March – May 2020. We sequenced samples from two groups: inpatients who were part of an observational study of COVID-19 in hospitalized individuals (n = 190), and symptomatic employees who presented to occupational health services (n = 135). All employees were diagnosed and treated in outpatient settings, except for one who was admitted as an inpatient. Genome copy number determined by qPCR of the nucleocapsid gene was highly variable and decreased by day from symptom onset (p < 0.001, linear model, Fig. 1A). We obtained 212 complete genomes (Fig. 1B), mostly from samples with higher viral loads (Fig. 1B). Consensus genomes had a median of 7 substitutions relative to the Wuhan-Hu-1/2019 reference sequence (range 4 – 12). Phylogenetic analysis of whole consensus genomes identified 10 unique evolutionary lineages in our cohort (lineages determined by the PANGOLIN system, see Methods; Fig. 1C). Most sequenced genomes fell in lineage B.1. We evaluated whether any employees were part of an epidemiologically linked cluster based on illness onset date, positive test status, and work location. We found that some employees were part of epidemiologically linked clusters (Fig. 1C). The genomes from clusters 2, 10, 19, 20, and one pair in cluster 29 had ≤ 1 consensus difference, while the rest had 2 – 7 differences. Many inter-cluster employee pairs also had identical or nearly identical consensus genomes. We have no information on epidemiologic linkage for the remaining sequenced individuals.

**Figure 1.**
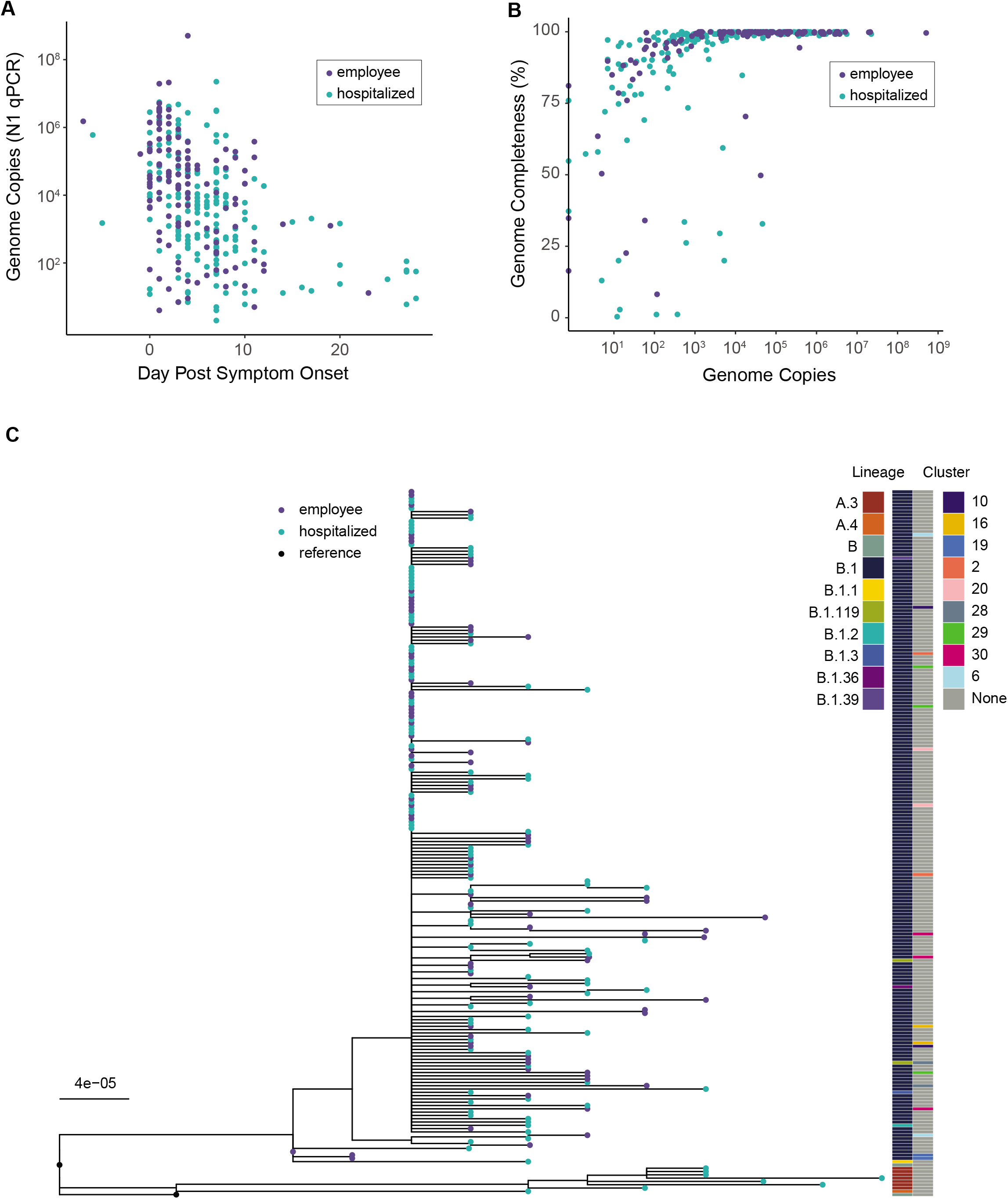
Viral shedding and overview of genome sequencing data. (A) Viral load by day of infection in hospitalized patients (teal) and employees (violet). Viral load, measured by qPCR of the N gene in units of genome copies per microliter of extracted RNA, is on the y-axis and day post symptom onset is on the x-axis. (B) Genome completeness by viral load in hospitalized patients (teal) and employees (violet). Viral load as shown in (A) is on the x-axis and the fraction of the genome covered above 10x read depth is shown on the y-axis. (C) Maximum-likelihood phylogenetic tree. Tips represent complete consensus genomes from hospitalized patients (teal) and employees (violet). The axis shows divergence from the root (Wuhan-Hu-1/2019). Heatmaps show PANGOLIN evolutionary linage (left) and epidemiologic cluster (right).

Identification of viral within-host variants can be prone to errors^22,23^. Therefore, we performed a mixing study to evaluate the accuracy of our pipeline for identifying intrahost single nucleotide variants (iSNV). We mixed two synthetic RNA controls that differ by seven single nucleotide substitutions at defined frequencies and input concentrations (Fig. 2A). These mixtures were sequenced using the same approach as the clinical samples. We identified true iSNV at the expected frequencies at ≥ 10^3^ copies/μL (Fig. 2B). There was greater variance in the observed variant frequencies at 10^2^ copies/μL compared to higher input concentrations. We obtained high sensitivity for iSNV at ≥ 2% frequency and ≥ 10^3^ copies/μL with sufficient genome coverage. Many false positive iSNV remained at ≥ 2% frequency and 10^2^ copies/μL despite multiple quality filters (Figure 2C, Supplemental Figure 1). However, false positive iSNV per sample drastically decreased with input concentrations ≥ 10^3^ copies/μL. Three false positive variants were identified in multiple samples above 10^4^ copies/μL: A3350U, G6669A, and U13248A. Because these iSNV were not randomly dispersed across the genome and were otherwise well-supported in the sequence data, we suspect that they represent low-frequency variants present in the synthetic RNA controls. Together, these data indicate that sufficient input viral load is a critical factor for accurate identification of iSNV.

**Figure 2.**
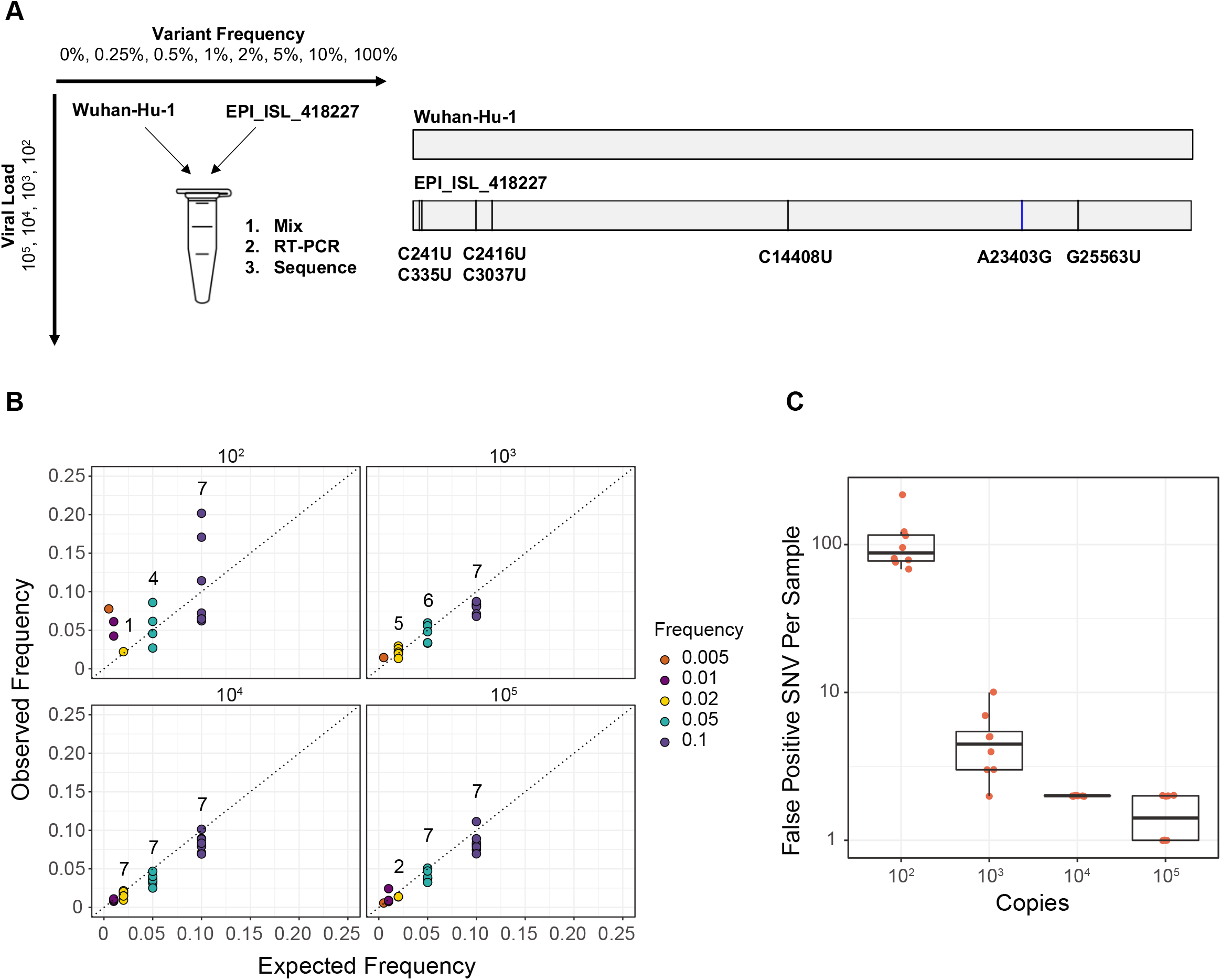
Assessing accuracy of intrahost variant detection by sequencing defined viral mixtures. (A) Schematic of the experiment. Wuhan-Hu-1 (reference) and EPI_ISL_418227 (variant) RNA were mixed at the given frequencies and viral loads (units of genome copies per microliter, representing the resulting mixture). Mixtures of RNA were amplified and sequenced in the same fashion as the clinical specimens. Reference and variant genomes differ by seven single nucleotide substitutions. (B) Observed frequency by expected frequency. Observed frequency of the true positive intrahost single nucleotide variants (iSNV) is on the y-axis and expected iSNV frequency is on the x-axis. Synthetic RNA copy number in units of genome copies per microliter of RNA is shown above each facet. Values above the points indicate the number of variants detected in that group (maximum of seven per group). (C) False positive iSNV. Number of false positive iSNV per sample is shown on the y-axis (base 10 log scale) and viral load as shown in (B) is on the x-axis. Each point represents a unique sample and the boxplots represent the median and 25^th^ and 75^th^ percentiles, with whiskers extending to the most extreme point within the range of the median ± 1.5 times the interquartile range.

Based on our benchmarking experiment, we identified iSNV in 178 specimens with viral loads ≥10^3^ copies/μL (Fig. 3A). We excluded position 11083, which is near a natural poly-U site and prone to sequencing errors^27^. Most specimens exhibited fewer than ten minor iSNV (median 1, IQR 0 – 3, Fig. 3B). There were four outlier specimens with greater than 15 iSNV. In these samples, iSNV were dispersed throughout the genome at various frequencies, so it is difficult to determine whether they represent mixed infections^11^. The locations of these samples on sequencing plates were not suggestive of cross-contamination. There was no difference in minor iSNV richness between hospitalized patients and employees treated as outpatients (p = 0.29, Mann-Whitney U test, Supplemental Figure 2). We identified more minor iSNV encoding non-synonymous changes than synonymous ones across most open reading frames (Fig. 3C) and identified more iSNV at lower frequencies (Fig. 3D), which together is suggestive of mild within-host purifying selection. Sample iSNV richness decreased with higher viral loads by about 1 iSNV per 10-fold increase in viral load (p = 0.01, multiple linear model, Supplemental Figure 3). Sample iSNV richness did not correlate with day from symptom onset (p = 0.75, multiple linear model, Fig. 3E). These results show that within-host diversity is low and remains that way over the duration of most SARS-CoV-2 infections.

**Figure 3.**
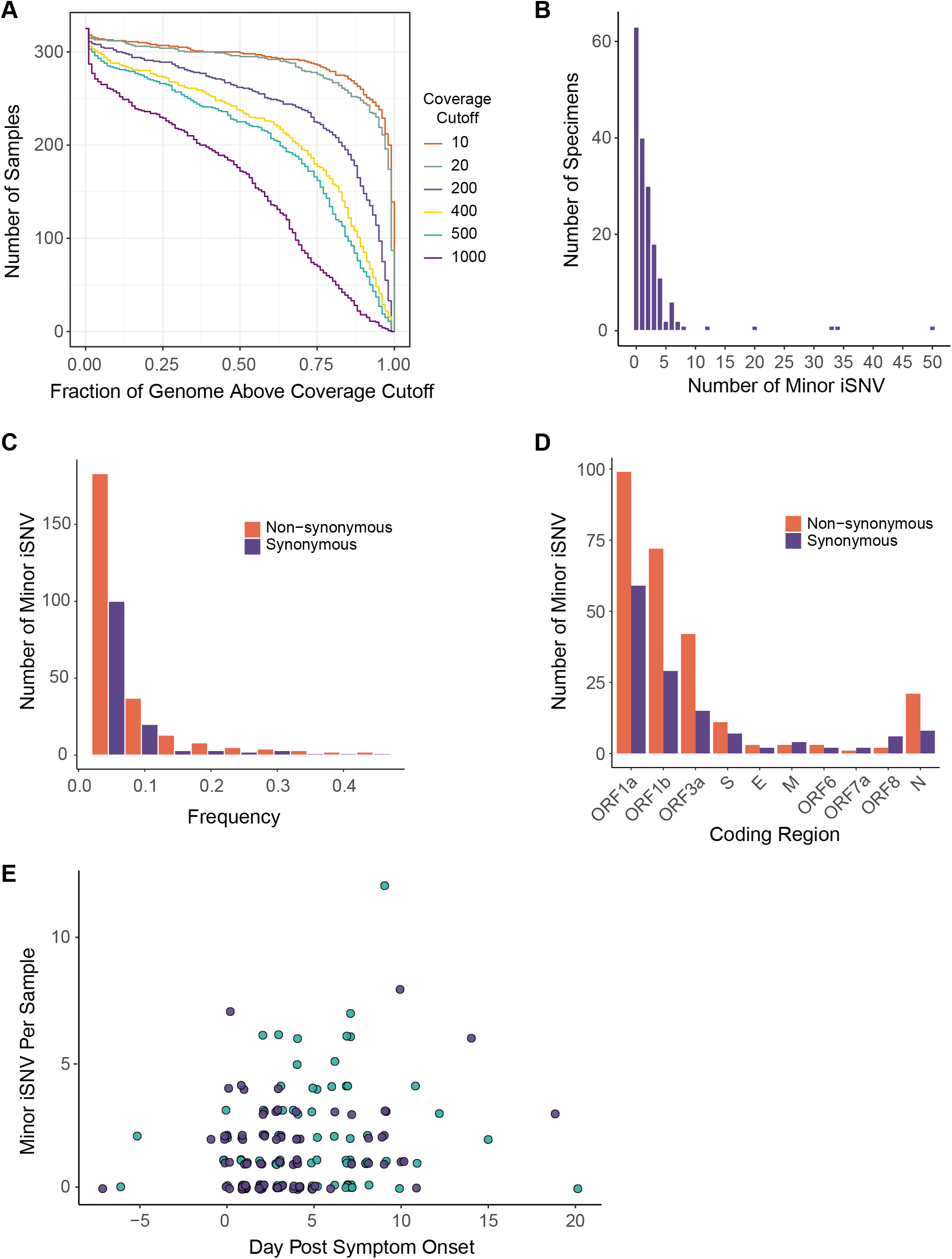
SARS-CoV-2 intrahost single nucleotide variant (iSNV) diversity. (A) Sequencing coverage for clinical samples. The number of clinical samples (y-axis) is shown by the fraction of the genome above a given read depth threshold (x-axis). The different lines show the data evaluated with six read depth thresholds. (B) Histogram of the number of specimens (y-axis) by the number of minor iSNV per sample (x-axis), n = 178. (C) Number of minor iSNV by frequency with a bin width of 0.05. Non-synonymous iSNV are shown in orange and synonymous iSNV are shown in violet. (D) Number of minor iSNV by coding region. Non-synonymous iSNV are shown in orange and synonymous iSNV are shown in violet. (E) Scatterplot of the number of minor iSNV per sample (y-axis) by the day post symptom onset (x-axis). Hospitalized patients are shown in teal and employees shown in violet. The four samples with > 15 iSNV shown in (B) are excluded from the plot for visualization.

Next, we investigated patterns of shared intrahost diversity between individuals. Most iSNV were unique to a single individual. However, 19 iSNV were present in multiple specimens (Fig. 4A). These did not include the three recurrent false positives found in the synthetic RNA controls. None of these mutations were located at sites known to commonly produce errors or homoplasies^27,28^. Two iSNV were present in three individuals (G12331A and A11782G, both synonymous changes in ORF1a) and one iSNV was present in six individuals (U13914G, encoding N149K in ORF1b). There was no clear phylogenetic clustering of genomes exhibiting these shared iSNV (Supplementary Figure 4). The U13914G mutation was shared between several sample pairs separated by 2 or more substitutions, and G12331A was shared between samples from different viral lineages (13 substitutions). These three mutations were first detected in our samples in late March 2020 (Fig. 4B). None reached > 1% frequency per week in consensus sequences submitted to GISAID through mid-November 2020. These results suggest that iSNV that arise convergently across viral lineages are not necessarily predictive of subsequent global spread of those mutations.

**Figure 4.**
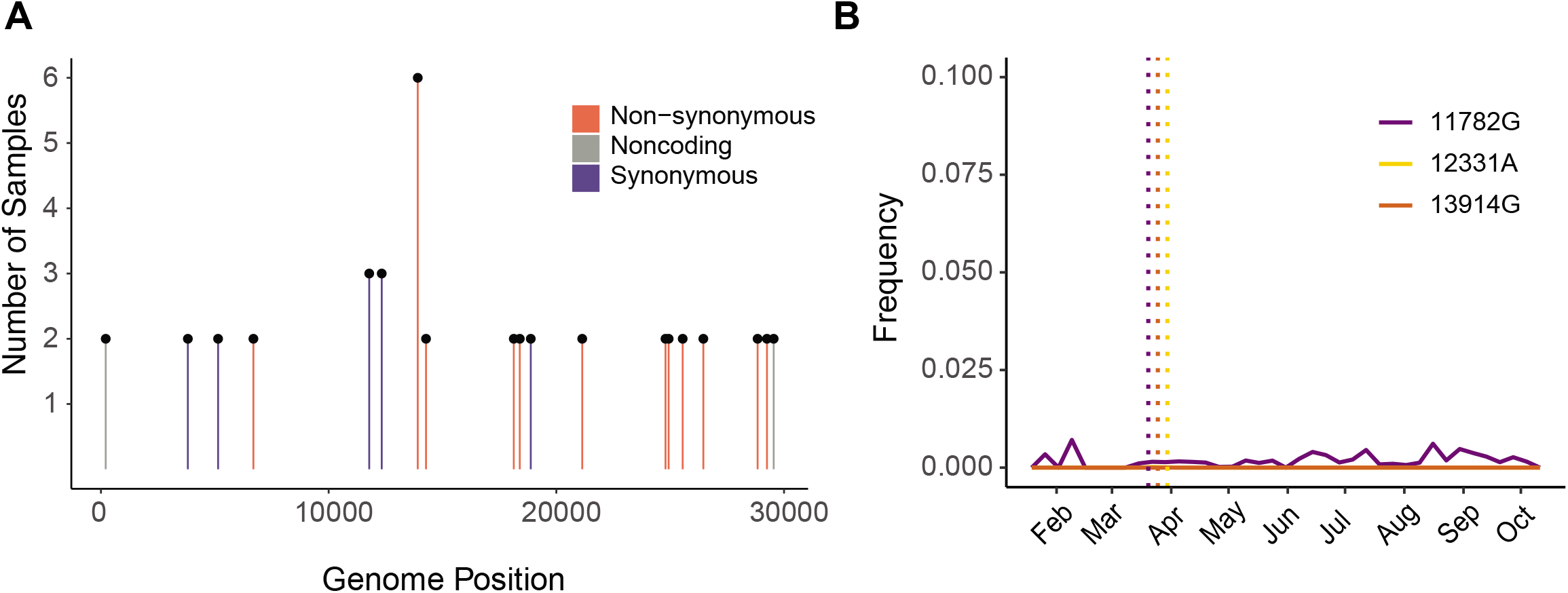
Shared iSNV across samples and their frequency in global consensus genomes. (A) Shared iSNV across samples, with the number of samples sharing the iSNV (y-axis) by the genome position (x-axis). Colors indicate the iSNV coding change relative to the reference. (B) The frequency (y-axis) of three iSNV shared by three or more samples over time (x-axis). The consensus genomes are from GISAID, as available on 2020-11-11. The vertical dotted lines represent the earliest time we detected each iSNV in our samples.

Transmission inference based on shared iSNV integrates information such as consensus genome sequences, sample dates, and shared iSNV^15^. Therefore, we compared shared iSNV across all unique pairs of specimens used for variant calling (n = 15753, Fig. 5). Because most iSNV were unique to an individual, most pairs did not share iSNV and only 0.23% of pairs shared one iSNV. Many pairs with shared iSNV were sequenced in separate batches, which reduces the likelihood that shared iSNV are due to cross-contamination. No employee pairs in the same epidemiologic cluster shared iSNV (see Fig. 1C). We identified fourteen unique pairs with shared iSNV between genomes that were near-identical (0 – 1 consensus differences), eight of which were collected within one week of each other. However, we have no epidemiologic data to suggest that these pairs of individuals are linked by transmission. We also identified shared iSNV between 23 pairs separated by ≥ 2 consensus substitutions (Fig. 5A and 5B) and 15 pairs with collection dates 7 – 28 days apart (Fig. 5B). Due to differences in viral lineage and time of collection, these are very unlikely to be transmission pairs. Together, these data indicate that iSNV can arise convergently between individuals who are unlikely to be related by transmission.

**Figure 5.**
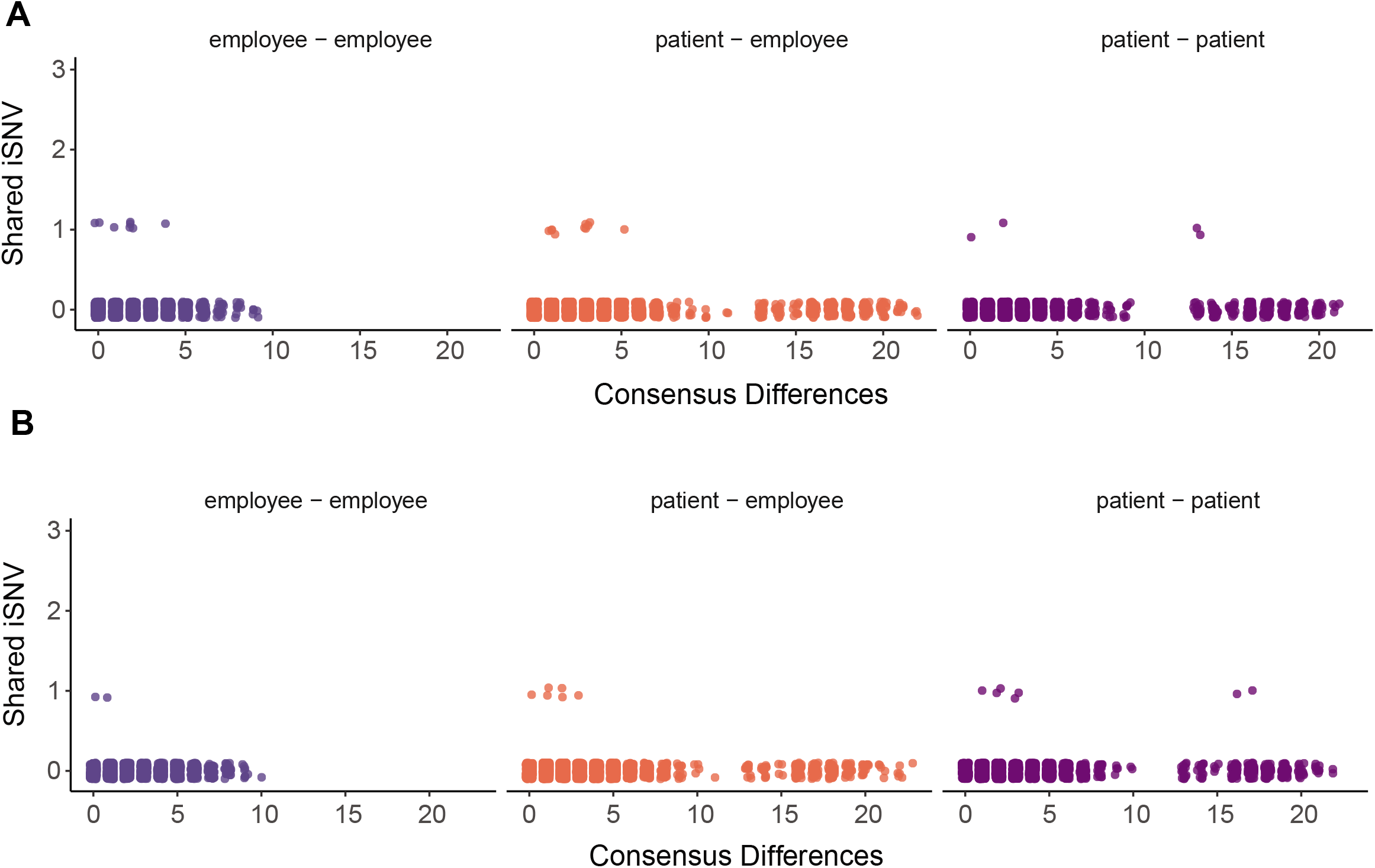
Pairwise comparisons of shared iSNV. Each unique pair is shown as a single point, with employee-employee pairs in violet (left), patient-employee pairs in orange (middle), and patient-patient pairs in purple (right). The number of iSNV shared by each pair is shown on the y-axis with the number of consensus differences between the pair of genomes on the x-axis. Pairs of samples collected within seven days of each other are displayed in (A), and pairs of samples collected greater than seven days apart are shown in (B).

## Discussion

Accurate characterization of SARS-CoV-2 intrahost diversity is important for understanding the spread of new genetic variants and its potential use in transmission inference. In this study, we sequenced upper respiratory specimens from a cohort of hospitalized COVID-19 patients and infected employees. We found that intrahost diversity is low and its distribution does not vary by time since symptom onset. We identified iSNV shared across viral genomes separated by time and disparate evolutionary lineages, indicating that iSNV can arise convergently. Because variants may be shared through parallel mutation rather than transmission, caution is warranted in the use of shared iSNV alone for inferring transmission chains. Intrahost variants shared across multiple individuals did not precede an increase in frequency in global consensus genomes, which suggests that identifying convergent iSNV may have limited utility in tracking broader SARS-CoV-2 evolution.

Specimen viral load is important when measuring intrahost diversity. We and others have shown that samples with low viral loads are prone to false positive iSNV and lower sensitivity^22,23,29^. A strength of our study is that we experimentally validated the accuracy of our variant calling by sequencing defined populations. Based on these results, we excluded samples with low viral load from subsequent analyses. Future studies of SARS-CoV-2 intrahost diversity should report and account for specimen viral loads to avoid this common source of error. We did not benchmark our sequencing approach for detecting insertions and deletions (indels) and therefore did not report these for the clinical specimens. Intrahost indels could conceivably provide useful information about within-host evolution, but accurate detection is also subject to similar issues of sample quality and viral load.

The low level of intrahost diversity that we found here is consistent with a recent preprint by Lythgoe et al.^9^. The fact that our work and the study by Lythgoe et al. were performed with different geographical areas, sequencing approaches (ARTIC Network amplicons vs. veSEQ metagenomic sequencing), and analysis methods lends credence to the results. Lythgoe et al. reported more shared variation than seen here, but this is most likely due to sequencing a greater number of samples among individuals within known epidemiologic clusters. We and Lythgoe et al. measure a lower level of intrahost diversity at the 2% frequency threshold compared to a recent study in Austria^12^. The reasons for this are not clear, but it is likely due to differences in sample viral loads and variant calling methods. We did not find a difference in intrahost diversity between hospitalized COVID-19 patients and those treated as outpatients, which suggests that viral diversity may not be a reliable marker for disease severity.

Measuring viral diversity over the course of infection is relevant for understanding how variants are transmitted to new hosts. Only genetic variants present at the time of a transmission event will have the opportunity to spread. Because SARS-CoV-2 usually transmits just before or several days after symptom onset^30,31^, it is important to define viral diversity in this window. Our cross-sectional analysis of diversity by time since symptom onset indicates that diversity does not significantly increase over the course of infection. A significant fraction of samples may not exhibit any iSNV at the time of transmission, which could limit the utility of iSNV for linking transmission pairs. Only a large bottleneck would lead to onward spread of most iSNV present during early infection. However, it is important to recognize that although the absolute level of diversity may not change over time, different variants may arise or go extinct during a given infection. This phenomenon was observed in a recent study by Tonkin-Hill et al.^11^. Serial samples from individuals could address this issue with higher resolution. Low diversity within hosts also shapes our expectations for emergence of resistance to drugs and monoclonal antibodies. With such limited substrate for selection to act upon, the short window of time between treatment and transmission could limit the spread of a variant selected within a host. Even during prolonged infections in immunocompromised hosts, there is only limited evidence of resistance to various COVID-19 therapeutics^32–34^.

Parallel evolution is a critical factor to consider in the interpretation of shared intrahost variation^15^. Even if iSNV identification were perfectly specific, iSNV can arise in parallel due to biological processes such as natural selection and genetic drift. A key finding of this work is that iSNV can arise in genomes that are unrelated by local transmission, specifically those across large time intervals and lineages. Shared iSNV between individuals with identical genomes collected the same week may also have arisen in parallel. These pairs are most likely not epidemiologically linked, but we are unable to rule out coincident local transmission in the community. Because iSNV can arise in parallel in genomes that are not linked by transmission, caution is needed when relying entirely on shared iSNV for transmission inference^11,13^.

We also found that identifying iSNV across multiple individuals did not precede an increase of those mutations in frequency in global consensus genomes. It is unclear whether these mutations arose due to positive selection, chance, or mutational “hotspots”^11^. It is possible that these mutations were lost due to purifying selection within hosts or during transmission^8,35^. These results suggest that iSNV may have lower utility for tracking broader SARS-CoV-2 evolution, but larger sample sizes in more geographic areas are necessary to evaluate this.

One of the most important variables for transmission inferences is the size of the transmission bottleneck^15^. If parallel evolution of iSNV occurs regularly and the transmission bottleneck is very small, that would increase the likelihood that shared iSNV are due to convergence rather than transmission. However, if the bottleneck is large, then iSNV may become more valuable for detecting transmission networks when consensus genomes are limited. There are currently conflicting results on the SARS-CoV-2 bottleneck size. Popa et al. estimated a bottleneck size of greater than 1000^12^. In contrast, Lythgoe et al. estimated a bottleneck size range from 1 – 8 based on 14 household pairs^9^. Lythgoe et al. in particular used extensive controls and validation for preventing contamination and identifying sequencing errors. Other studies both in humans and in domestic cats have estimated small bottlenecks^36,37^. It is difficult to interpret these contrasting results because each study used different sequencing and analysis methodologies. In recent work on influenza A virus, a study of methodological differences was key for resolving different conclusions about the bottleneck size^38^. One factor that has not yet been clearly defined is how the time interval between donor-recipient pairs affects SARS-CoV-2 bottleneck estimates. We expect that further work will clarify the reasons behind these conflicting estimates.

Because of the high incidence and low mutation rate of SARS-CoV-2, genomic epidemiology is necessarily constrained in its ability to determine exact transmission chains in an outbreak. Using minor genetic variation to increase the resolution of genomic epidemiology requires attention to the underlying processes of within-host viral evolution and awareness of possible confounders. Unified statistical frameworks that incorporate sequences, metadata, and epidemiological models are likely the most robust approaches for integrating intrahost variants, but these models also must account for parallel evolution^15–17^. As others have recently suggested^11^, we caution against assigning transmission pairs solely by virtue of shared iSNV in the absence of clear epidemiologic information.

## Acknowledgements

We thank the University of Michigan Clinical Microbiology Laboratory and the University of Michigan Central Biorepository for their assistance in providing samples. We thank Christina Cartaciano and the University of Michigan Microbiome Core for their assistance in sequencing. We thank Emily Stoneman from Michigan Medicine Occupational Health Services for assistance with employee data. This work was supported by a University of Michigan COVID-19 Response Innovation Grant (to ASL), K01AI141579 (to JGP) and CDC U01 IP000974 (to ETM)

## Materials and Methods

We collected clinical metadata and residual diagnostic specimens positive for SARS-CoV-2 from hospitalized patients enrolled in the CDC HAIVEN (Hospitalized Adult Influenza Vaccine Effectiveness Network) study and infected employees enrolled in the HARVI (hospital associated respiratory virus infection) study. These studies and the use of residual specimens were approved by the University of Michigan Institutional Review Board.

Date of illness onset for hospitalized patients was collected individually via medical chart abstraction from physician notes. Michigan Medicine employees with any suspected COVID-19 symptoms were asked to call a COVID-19 healthcare worker hotline before reporting to work. Date of symptom onset, a list of symptoms, close contacts, travel history, and work location and description were recorded. After testing, employee clusters were determined by illness onset date, positive test status, and work location.

### Genome amplification and sequencing

Residual samples from nasopharyngeal swabs and sputum specimens were centrifuged at 1200 x g. and 200 microliters were aliquoted. RNA was extracted with the Invitrogen PureLink Pro 96 Viral RNA/DNA Purification Kit and eluted in volumes of 100 microliters. Complementary DNA was reverse transcribed with SuperScript IV (ThermoFisher). The SARS-CoV-2 genome was amplified in two multiplex PCR reactions using the ARTIC Network V3 primer sets. Sequencing libraries were prepared with the NEBNext Ultra II kit and pooled in equal volumes after barcoding. The pooled sequencing library was gel extracted to remove adapter dimers. Libraries were sequenced on an Illumina MiSeq at the University of Michigan Microbiome Core facility (v2 chemistry, 2×250 cycles). To validate this approach, we used two synthetic RNA controls that differ by seven single nucleotide mutations, Wuhan-Hu-1 and EPI_ISL_418227 (Twist Bioscience, San Francisco, CA). We mixed the two RNAs at various copy numbers (10^5^, 10^4^, 10^3^, 10^2^ genome copies/μL) and frequencies (0%, 0.25%, 0.5%, 1%, 2%, 5%, 10%, and 100%). We amplified and sequenced each RNA mixture as described above.

### Viral load measurements

We measured SARS-CoV-2 genome copy concentration for each sample by qPCR using conditions outlined in the CDC 2019-Novel Coronavirus EUA protocol (https://www.fda.gov/media/134922/download). The nucleocapsid gene was amplified using the CDC N1 primer and probe set as follows: 2019-nCoV_N1 Forward Primer GACCCCAAAATCAGCGAAAT; 2019-nCoV_N1 Reverse Primer TCTGGTTACTGCCAGTTGAATCTG; 2019-nCoV_N1 Probe ACCCCGCATTACGTTTGGTGGACC. Probe sequences were FAM labeled with Iowa Black quencher (Integrated DNA Technologies, Coralville, IA). Reactions were performed using TaqPath 1-step RT-qPCR master mix (Thermofisher, Waltham, MA) with 500 nM of each primer and 250 nM of each probe in a total reaction volume of 20 µl. Cycling conditions were as follows: 2 min at 25 °C, 15 min at 50 °C, 2 min at 95 °C, and 45 cycles of 3 seconds at 95 °C, 30 seconds at 55 °C. Samples were run on an Applied Biosystems 7500 FAST real-time PCR system. Cycle threshold (Ct) was designated uniformly across PCR runs. Standard curves based on serial dilutions of a plasmid containing the nucleocapsid sequence were used to determine copy number for each plate of samples. Copy number is expressed in genome copies per microliter of extracted viral RNA.

### Analysis of sequence reads

We aligned reads to the MN908947.3 reference genome with BWA-MEM version 0.7.15^39^. We removed sequencing adaptors and trimmed ARTIC primer sequences with iVar 1.2.1^23^. We determined the consensus sequences with iVar 1.2.1, taking the most common base as the consensus (>50% frequency). We placed an N at positions along the MN908947.3 reference with fewer than 10 reads. We manually inspected insertions and deletions by visualizing alignments with IGV (version 2.8.0)^40^. We identified single nucleotide variants with iVar 1.2.1 using the following parameters: sample with viral load ≥ 10^3^ copies/μL; sample with consensus genome length of ≥ 29000; sample with ≥ 80% of genome sites above 200x coverage; iSNV frequency threshold of 2%; read depth of ≥ 100 at iSNV sites; ≥ 10 reads with average Phred score of > 35 supporting a given iSNV; iVar p-value of < 0.0001. All samples on which we called variants had > 50,000 mapped reads. We accounted for strand bias by performing a two-sided Fisher’s exact test for hypothesis that the forward/reverse strand counts supporting the variant base are derived from the same distribution as the consensus base. We then applied a Bonferroni multiple test correction and excluded variants with an adjusted p-value < 0.05. To generate a phylogenetic tree, we aligned consensus genomes with MUSCLE 3.8.31 and masked positions that are known to commonly exhibit homoplasies or sequencing errors^41^. We generated a maximum likelihood phylogeny with IQ-TREE, using a GTR model and 1000 ultrafast bootstrap replicates^42,43^. Evolutionary lineages (Pango lineages) were assigned with PANGOLIN^44^.

### Data and code availability

Raw sequence reads are available as fastq files from the Sequence Read Archive at accession number PRJNA682212, with human-mapping reads removed. Analysis code is available at https://github.com/lauringlab/SARSCov2_Intrahost. Consensus genome sequences are publicly available at the GitHub link and on GISAID.

## Supplemental Figure Legends

**Supplemental Figure 1**. True and false positive iSNV in RNA mixture validation experiment. Each iSNV is shown as a point, with the frequency on the y-axis and genome position on the x-axis. True positive iSNV are shown in violet and false positive iSNV are shown in orange. All iSNV displayed have a frequency of 2% or greater. Viral loads are shown above each facet, in units of genome copies per microliter of RNA.

**Supplemental Figure 2**. Number of minor iSNV per sample (y-axis) across groups, with hospitalized patients shown by teal points and employees shown by violet points. Boxplots for each group represent the median and 25^th^ and 75^th^ percentiles, with whiskers extending to the most extreme point within the range of the median ± 1.5 times the interquartile range.

**Supplemental Figure 3**. Number of minor iSNV per sample (y-axis) by genome copies per microliter of RNA (x-axis). Hospitalized patients are shown by teal points and employees shown by violet points.

**Supplemental Figure 4**. Maximum likelihood phylogenetic tree as shown in Figure 1C. Tips represent complete consensus genomes from hospitalized patients (teal) and employees (violet). The x-axis shows divergence from the root (Wuhan-Hu-1/2019). Heatmaps show samples that contain each of the three mutations as an iSNV.

