## Supplemental Figure 1 for "Temporal dynamics of SARS-CoV-2 mutation accumulation within and across infected hosts"

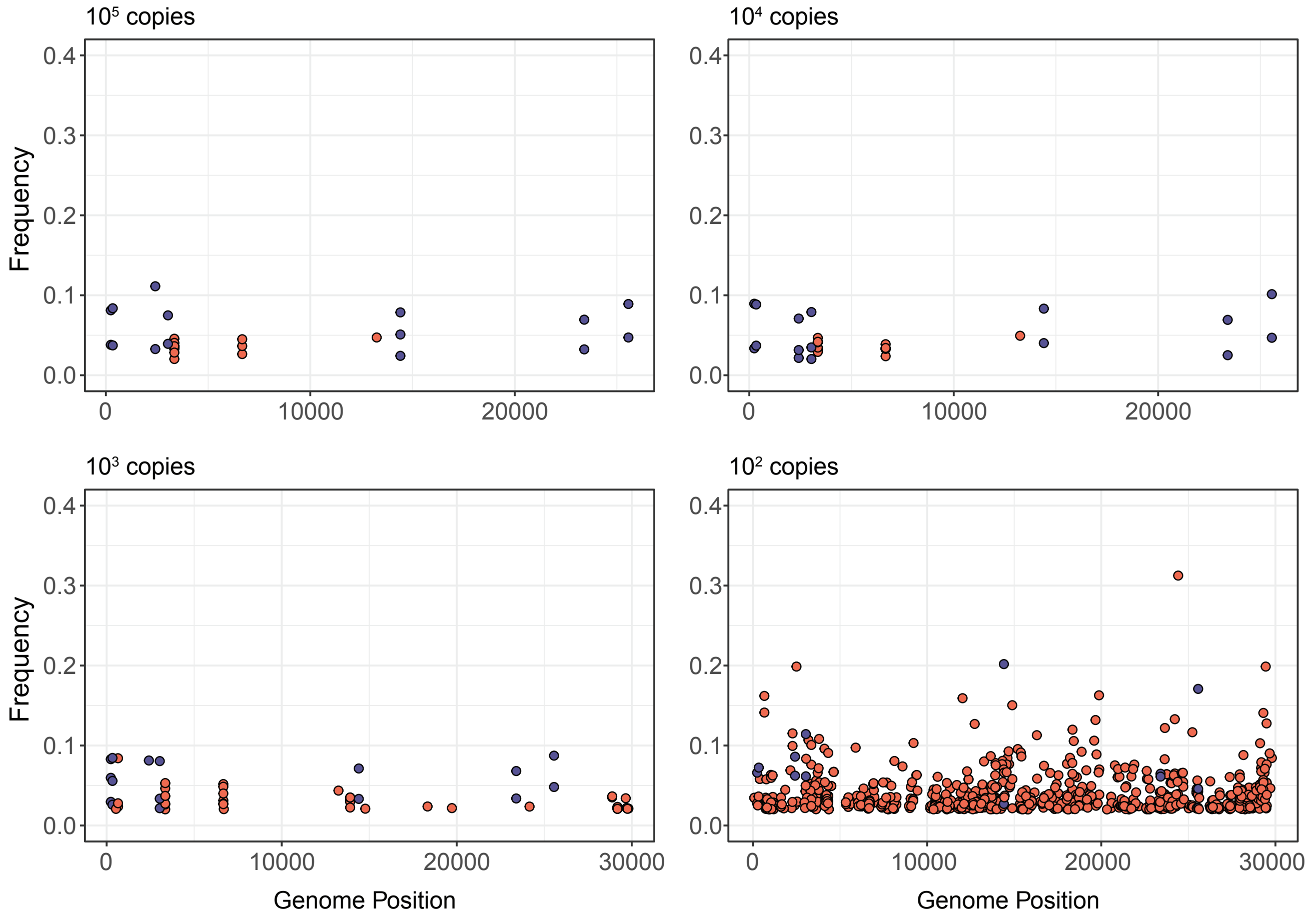


Supplemental Figure 1. True and false positive iSNV in RNA mixture validation experiment. Each iSNV is shown as a point, with the frequency on the y-axis and genome position on the x-axis. True positive iSNV are shown in violet and false positive iSNV are shown in orange. All iSNV displayed have a frequency of 2% or greater. Viral loads are shown above each facet, in units of genome copies per microliter of RNA.
