## Supplemental Figure 2 for "Temporal dynamics of SARS-CoV-2 mutation accumulation within and across infected hosts"

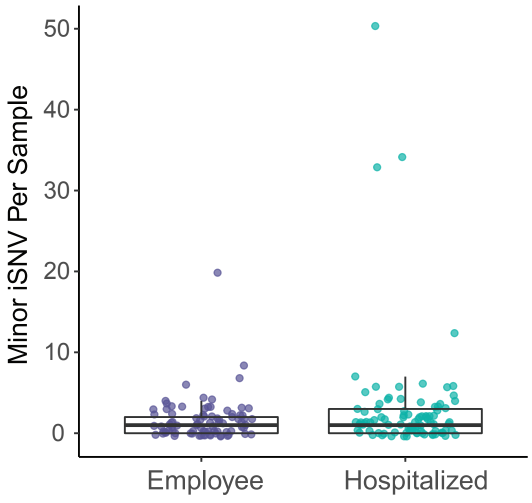


Supplemental Figure 2. Number of minor iSNV per sample (y-axis) across groups, with hospitalized patients shown by teal points and employees shown by violet points. Boxplots for each group represent the median and 25th and 75th percentiles, with whiskers extending to the most extreme point within the range of the median ± 1.5 times the interquartile range.
