## Supplemental Figure 3 for "Temporal dynamics of SARS-CoV-2 mutation accumulation within and across infected hosts"

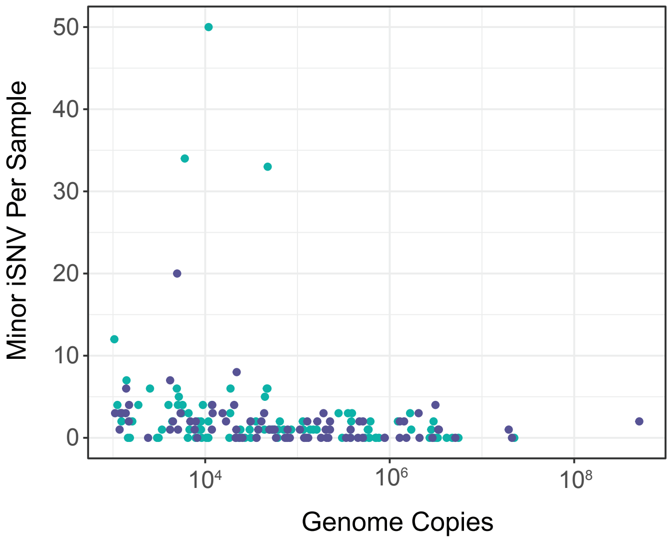


Supplemental Figure 3. Number of minor iSNV per sample (y-axis) by genome copies per microliter of RNA (x-axis). Hospitalized patients are shown by teal points and employees shown by violet points.
