## Supplemental Figure 4 for "Temporal dynamics of SARS-CoV-2 mutation accumulation within and across infected hosts"

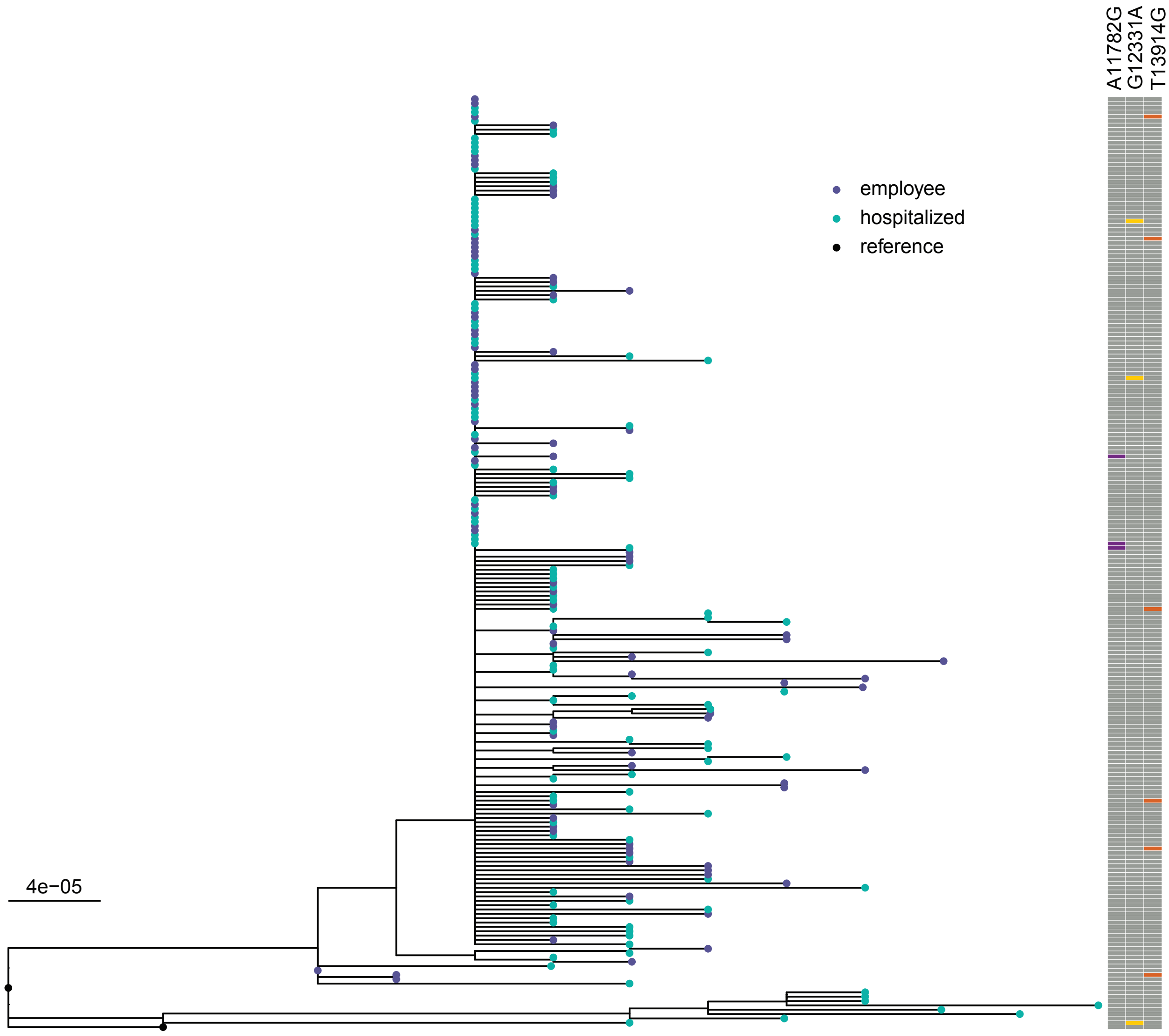


Supplemental Figure 4. Maximum likelihood phylogenetic tree as shown in Figure 1C. Tips represent complete consensus genomes from hospitalized patients (teal) and employees (violet). The x-axis shows divergence from the root (Wuhan-Hu-1/2019). Heatmaps show samples that contain each of the three mutations as an iSNV.
